# Heatwaves and cold snaps alter host-parasite population dynamics in the *Daphnia magna-Ordospora colligata* system

**DOI:** 10.1101/2025.08.01.668077

**Authors:** Niamh McCartan, Louise Bezborodko, Joseph Strawbridge, Floriane O’Keeffe, Sadie DiCarlo, Pepijn Luijckx

## Abstract

Climate change is driving more frequent and severe temperature extremes, including heatwaves and cold snaps, with growing implications for ecology and disease. Yet, our understanding of how heatwaves and cold snaps influence disease dynamics remains underexplored. Using the host *Daphnia magna* infected with its microsporidian microparasite *Ordospora colligata,* pathogen fitness and host population size were measured in experimental populations using a factorial design at four baseline temperatures (14, 17, 20 and 23°C). A heatwave or cold snap treatment with an amplitude of ±6°C was administered four weeks after measurements began and lasted for ten days. The effect of heatwaves is dependent on baseline temperature but can induce long-lasting increases in burden (>4 weeks). The impact of cold snaps were also temperature-dependent, leading to short-term increases in parasite fitness at higher temperatures. Host population size also varied in response to temperature and treatment. Importantly, burden and host density were interdependent, jointly shaping infection patterns. At lower temperatures, parasite burden and host population size were positively correlated, whereas at higher temperatures, increased host population size corresponded with reduced burden. These patterns were consistent at both individual and population levels, underscoring how individual physiological responses can scale up to impact disease dynamics across populations. Thus, extreme temperature variation can have complex, context-specific outcomes on disease dynamics. As climate extremes become more frequent, understanding these nuanced responses is critical for predicting and managing disease risk in natural populations.

**Author Summary:** We are experiencing more extreme weather events around the world, including heatwaves and cold snaps, but we don’t fully understand how these temperature extremes will affect wildlife diseases. In our study, we tested how heatwaves and cold snaps influence both parasite success and host population size using a small aquatic animal, the water flea, and its naturally occurring gut parasite. We ran experiments at four average temperatures and simulated a heatwave or cold snap by raising or lowering the temperature by 6°C for ten days. We found that heatwaves often led to long-lasting increases in parasite burden, while cold snaps caused short-term spikes in parasite fitness, especially at warmer average temperatures. These effects also depended on the baseline temperature and were linked to changes in the host population. At cooler temperatures, parasite levels increased as host populations grew, but at warmer temperatures, the opposite happened, resulting in negative density-dependence. This suggests that the impact of extreme weather on disease isn’t straightforward; it depends on when and where the event occurs. As extreme temperatures become more common with climate change, understanding these complex interactions is important for predicting disease outbreaks in the wild.

## Introduction

Climate change is transforming ecological and epidemiological systems by altering the environment and influencing interactions among organisms [1] with far-reaching consequences for human health [2], ecosystem functioning [3], and the global economy [4]. Changing global temperatures have shifted the geographic ranges of both free-living organisms [5] and pathogens [6], affected the frequency and severity of disease outbreaks [7], influenced host immunity [8] and resulted in higher disease prevalence in keystone species, leading to declines in biodiversity [9]. For example, a recent study projected that rising global temperatures will expand the geographic and seasonal range of mosquito-borne diseases like malaria and dengue, potentially placing an additional 4.7 billion people at risk of infection by 2070 [10]. Indeed, host-parasite relationships may be particularly vulnerable to temperature changes [11] as temperature can influence host-pathogen interactions in numerous and complex ways. For example, the effect of changing mean temperature can be direct by altering parasite development [12], transmission [13], and survival in the environment [14]. Temperature can also act indirectly by influencing host behaviour [15], immunity [8], or modification of physical barriers in the host [16]. Moreover, temperature-driven shifts in host mortality [17], fecundity [18], and feeding strategies [19] can reshape host and parasite population trajectories over time, potentially altering outbreak dynamics [20], population stability [21], and long-term persistence [11]. Not only do rising global temperatures affect many aspects of host-pathogen interactions simultaneously, but they can also have opposing effects on different traits. For example, McCartan et al. (2024) found that after a cold snap, infectivity decreased with increasing temperature while within-host growth of the pathogen accelerated. While the relationship between rising global mean temperature and disease is complex, extreme weather events and altered temperature variation associated with global warming may further complicate disease outcomes.

Climate change not only raises mean global temperatures but also increases the frequency, intensity, and unpredictability of extreme temperature events such as heatwaves and cold snaps [17]. These anomalous weather events are becoming more common and prolonged, with heatwaves accelerating in frequency and duration since the 1950s [18] and cold snap frequency accelerating since 1979 [19]. Such extreme temperature events can have pronounced effects on host-pathogen systems, as both heatwaves and cold snaps may challenge the physiological limits of hosts and parasites [20–22]. For example, simulated heatwaves facilitated the invasion of a lowland *Drosophila* species into high-elevation habitats, leading to cascading effects on resident parasitoid species and their host interactions [23]. Similarly, cold snaps can increase disease burden up to five-fold or decrease it by three-fold, depending on the baseline temperature [24]. Moreover, these temperature anomalies can destabilise population dynamics [25], alter epidemic cycles [26], or, in some cases, cause population crashes or regime shifts [27]. Yet, despite growing evidence of their ecological significance, the role of anomalous temperature events in shaping disease dynamics at the population level remains underexplored [25, 28], especially in systems where temperature variability interacts non-linearly with parasite transmission and host resilience.

Here, we examined the effect of heatwaves and cold snaps on disease dynamics in experimental populations of *Daphnia magna* infected with its microsporidium micro-parasite *Ordospora colligata* (a unicellular gut pathogen) across a broad temperature range. This study was based on previous work using the same host-pathogen system [20, 24, 29], which studied the effects of extreme weather events at an individual level. These studies found that even minor shifts in temperature lead to increased parasite performance, as suggested by the Thermal Variability Hypothesis [30]. Declines in performance under extreme thermal stress were also observed when temperatures exceeded their physiological limits, a pattern described by the Thermal Stress Hypothesis [31]. Moreover, these studies also showed differential thermal sensitivity of *Daphnia* and *Ordospora*, such that mismatches in performance optima favoured either the host or the parasite depending on environmental conditions, which conforms to the Thermal Mismatch Hypothesis [32]. These results indicate that temperature can profoundly and unpredictably alter infection outcomes at the individual level. These physiological and interaction-level responses can scale up to influence epidemic trajectories [33], drive changes in host density [34], and shift host-parasite distribution [35]. Yet, whether the previous observations at the individual level in the *Daphnia-Ordospora* model system scale up to the population level remains an open question.

Here, we tested whether the effects observed at the individual level could inform population-level outcomes using populations of *Daphnia magna* infected with the microsporidium gut parasite *Ordospora colligata*. We compared exposed populations, in which *O. colligata* was well established and maintained at constant baseline temperatures, to those that experienced either a heatwave or a cold snap (5 replicate populations for all treatments). Lower baseline temperatures (14 and 17°C) experienced a +6°C heatwave, while higher baseline temperatures experienced a −6°C cold snap (20 and 23°C). Both heatwave and cold snap treatments lasted the same duration (10 days) and started at the same time (4 weeks after measurements began) (Fig 1). Measurements (population size and pathogen burden) for each population were recorded weekly until the end of the experiment (week 9). Population results were compared to previously published work that conducted comparable experiments on within host proliferation and infection. We expected that extreme temperature variation would generate complex, context-dependent outcomes in both parasite burden and population size, depending on the baseline temperature, with potential consequences for long-term population stability and disease dynamics under climate change.

**Fig 1:**
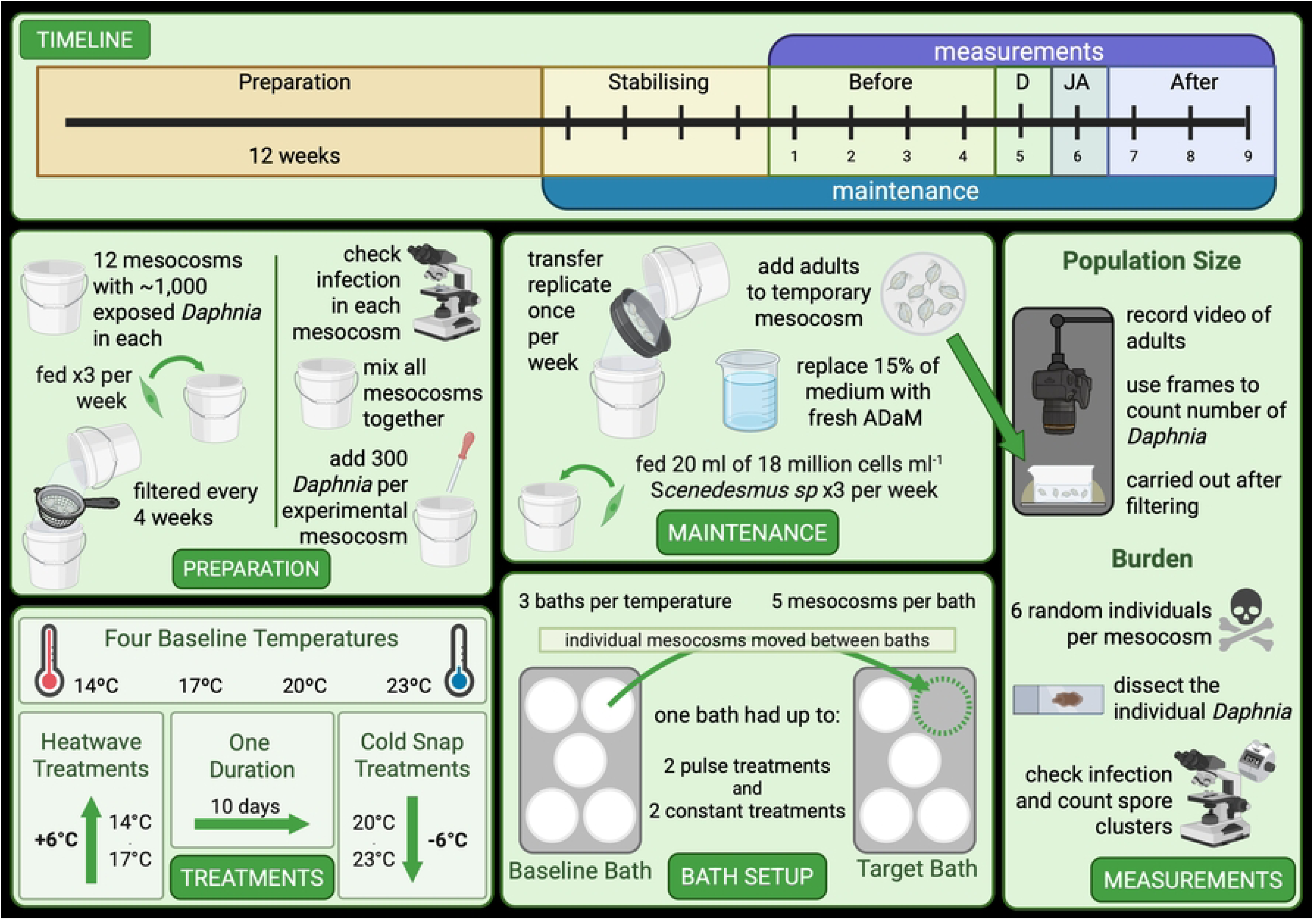
**Experimental Setup.** TIMELINE; illustrates the timeline with twelve weeks of preparation, four weeks of stabilising, and nine weeks of measurements. The pulse treatment began on week four, and measurements during the heatwave (D) and the cold snap occurred in week five. All pulse treatments ended after 10 days, so the counts occurring in week six were just after the pulse treatment (JA). PREPARATION; preparation started twelve weeks prior to exposure when ∼1,000 exposed Daphnia were added to 2 L mesocosms. The week before the experiment began, all mesocosms were checked for infections and then mixed. From this mix, 300 Daphnia of varying sizes were added to an experimental mesocosm. Replicate one began on day one, and then the next replicate began the day after to stagger maintenance and measurements. TREATMENTS; eight treatments were included in the experiment (four baseline temperatures • two pulse treatments [either heatwave or cold snap]), and all baseline temperatures also had constant controls. Each treatment had five replicates. BATH SETUP; each temperature had three baths, and each bath held five mesocosms. A bath contained up to two pulse treatments and two constant treatments. To simulate a pulse treatment, mesocosms were moved between baths and then returned to the baseline bath when the pulse treatment was finished. MAINTENANCE; maintenance was carried out once a week when the Daphnia were filtered to remove debris, and 15% of the medium was changed for fresh medium. MEASUREMENTS; adult Daphnia were separated from the rest of the population by sieving into a temporary mesocosm, and two videos were taken for population size analysis. On the same day, six Daphnia from the mesocosm were dissected to check for infection. Figure created on Biorender.com.

## Results

### Parasite fitness

The prevalence of infection was nearly 100% in all *Daphnia* populations, irrespective of the treatment or temperature regime (p-value not significant), although burden differed between treatments. Extreme temperature variation (that is, a pulse treatment) altered parasite proliferation up to 2.9-fold depending on the baseline temperature and timing in relation to the event (Fig 2). Specifically, the mean burden of the heatwave treatment was higher during a heatwave and remained higher for one week after, at a baseline temperature of 14°C (Fig 2A) *(518 vs 256 and 574 vs 198 spore clusters, respectively, emmean, z(inf) = −2.625, p = 0.041, and emmean, z(inf) = −3.989, p < 0.001*, *see Table 5S for p-values of all custom contrasts*). Moreover, although no discrete ‘after’ treatments were significant (i.e., no significant custom contrasts), thermal pulses resulted in long-term effects by shifting the trajectory of burden over time (*GLMM, Estimate = 0.173, p = 0.035, Table 1A*); specifically at 14°C, long-term effects were present after a heatwave occurred (*14°C GLMM, Estimate = 0.567, p = 0.007, Table 3SA*). Therefore, temperature modulated the persistence of pulse treatment effects, with diminished increases in burden at higher temperatures (*GLMM, Estimate = –8.22, p = 0.042, Table 1A*). That extreme temperature can influence the outcome of parasite fitness is also relevant to cold snaps, as the burden increased just after the cold snap occurred (week 6) at a baseline temperature of 23°C (409 vs 207 spore clusters) (Fig 2D) (*emmean, z(inf) = −2.562, p = 0.042, see Table 5S*). However, at baseline temperatures of 17 and 20°C, extreme temperature events did not alter the disease burden in the populations during any period (*no contrasts were significant; see Table 5S for a full list of contrasts*). Thus, not all extreme temperature events will lead to the same changes in disease outcomes, and the outcome is dependent on the baseline temperature and time in relation to the pulse event (*GLMM, Type III Wald χ²_(8)_ = 40.63, p < 0.001, Table 1B, Fig 2*). Together, these results demonstrate that the effects of thermal extremes on parasite burden are highly context-dependent and temporally dynamic, varying with both baseline temperature and the timing of the event, factors which may also interact with other ecological drivers such as host population dynamics.

**Fig 2:**
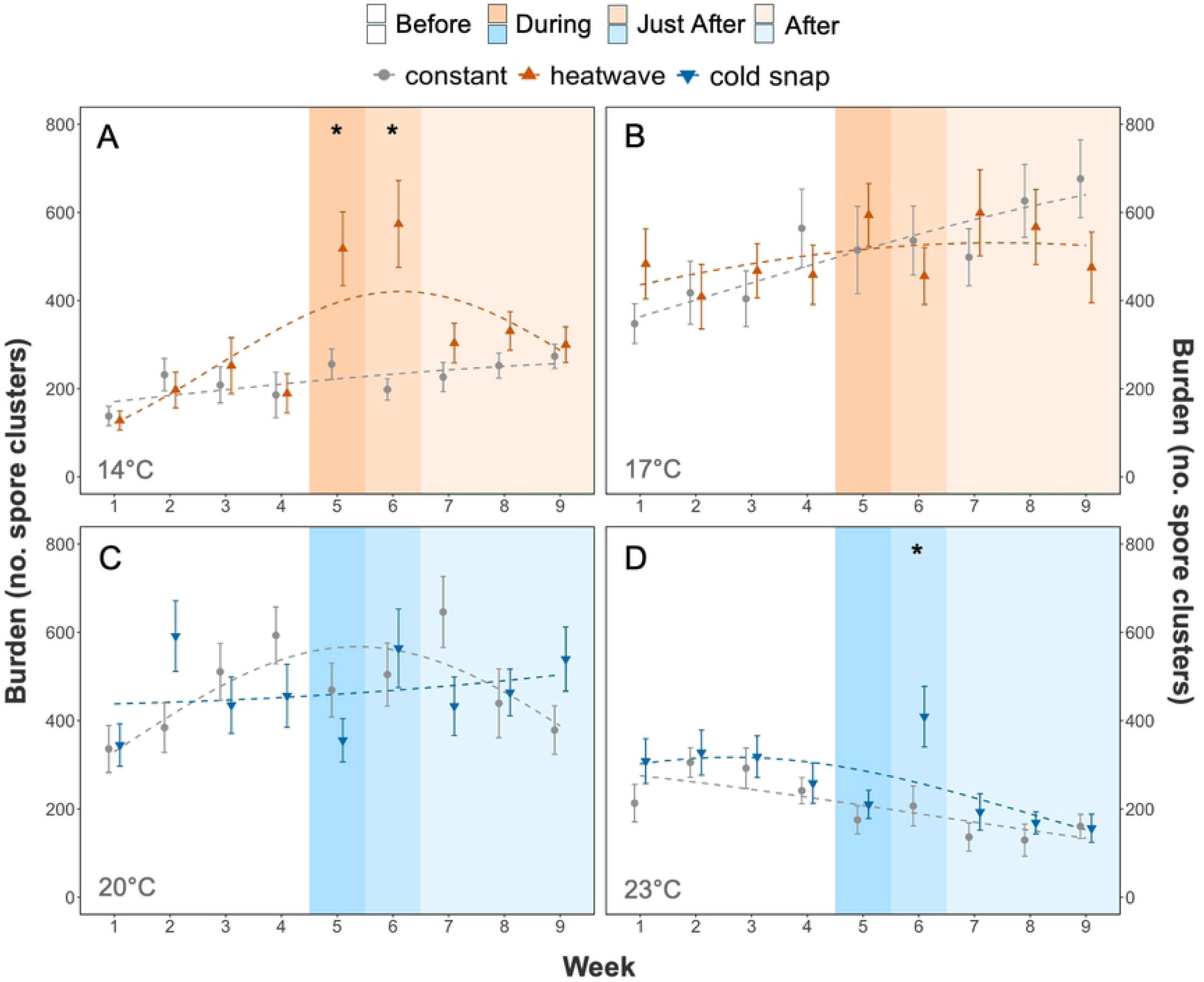
**The temporal effects of pulse temperature treatments on burden** in Daphnia magna infected with Ordospora colligata. Burden (mean number of spore clusters, including zeros for uninfected but exposed hosts) is shown over the nine recorded weeks at four baseline temperatures: (A) 14°C, (B) 17°C, (C) 20°C, and (D) 23°C. Each spore cluster contains 32-62 individual Ordospora. Error bars represent the standard error, and each dashed line is the quadratic fit of the generalised linear mixed model (see Table 1A for statistical results). Grey; constant treatment, orange; heatwave pulse treatment, blue; cold snap pulse treatment. Shaded regions mark experimental periods: Before (white), During (orange/blue), Just After (lighter orange/blue), and After (lightest orange/blue). Asterisks (*) indicate significant differences in burden between pulse and constant treatments in specific time periods (emmeans contrasts, Table 5S).

**Table 1:** **Type III Wald chi-square test results for the generalised linear mixed model (GLMM) analysing burden** (mean number of spore clusters, including zeros for uninfected but exposed hosts) in Daphnia magna infected with Ordospora colligata. A; GLMM, B; Type III Wald χ² test. The GLMM included fixed effects of temperature (modelled as a quadratic polynomial and separated by each degree, linear [L] and quadratic [Q]), time period (relative to a heatwave or cold snap), and population size, with a random effect of mesocosm. The Type III Wald χ² tests the significance of each predictor in the context of the full model. Significant values are bolded.

| **A** | **Explanatory Variable** | **Estimate** | **SE** | **z value** | **p value** |
| --- | --- | --- | --- | --- | --- |
|  | **Intercept** | 5.880 | 0.083 | 71.200 | <0.001 |
|  | Temperature (L) | 0.472 | 2.223 | 0.210 | 0.832 |
|  | **Temperature (Q)** | **-18.790** | **2.356** | **-7.970** | **<0.001** |
|  | Period before | -0.080 | 0.077 | -1.040 | 0.300 |
|  | **Period during** | **0.431** | **0.128** | **3.370** | **<0.001** |
|  | **Period just after** | **0.660** | **0.119** | **5.530** | **0.000** |
|  | **Period after** | **0.173** | **0.082** | **2.100** | **0.035** |
|  | **Population size** | **0.001** | **0.000** | **4.220** | **<0.001** |
|  | **Temperature (L): Period before** | **9.637** | **3.846** | **2.510** | **0.012** |
|  | Temperature (Q): Period before | 5.250 | 3.658 | 1.440 | 0.151 |
|  | **Temperature (L): Period during** | **-17.830** | **6.078** | **-2.930** | **0.003** |
|  | **Temperature (Q): Period during** | **13.800** | **5.958** | **2.320** | **0.021** |
|  | Temperature (L): Period just after | -3.625 | 5.455 | -0.660 | 0.506 |
|  | **Temperature (Q): Period just after** | **19.460** | **5.508** | **3.530** | **<0.001** |
|  | **Temperature (L): Period after** | **-8.220** | **4.049** | **-2.030** | **0.042** |
|  | Temperature (Q): Period after | 4.189 | 3.778 | 1.110 | 0.268 |
|  | **Temperature (L): Population size** | **-0.032** | **0.014** | **-2.300** | **0.021** |
|  | **Temperature (Q): Population size** | **-0.043** | **0.012** | **-3.480** | **0.001** |

| **B** | **Explanatory Variable** | **Chisq** | **Df** | **P value** |
| --- | --- | --- | --- | --- |
|  | **(Intercept)** | **5069.743** | **1** | **<0.001** |
|  | **Temperature** | **53.612** | **2** | **<0.001** |
|  | **Time Period** | **39.977** | **4** | **<0.001** |
|  | **Population** | **33.267** | **1** | **<0.001** |
|  | **Temperature: Time Period** | **40.634** | **8** | **<0.001** |
|  | **Temperature: Population** | **16.765** | **2** | **<0.001** |

Population size (i.e., the number of large reproducing adult females) was also important in predicting parasite fitness, both in relation to baseline temperature and independently of other influences (*GLMM, Type III Wald χ²_(2)_ = 16.77, p < 0.001, and GLMM, Type III Wald χ²_(1)_ = 39.98, p < 0.001, respectively, Table 1B*). While larger populations were generally associated with higher parasite burden (*17°C GLMM, Estimate = 0.002, p = 0.003, Table 3SB, Fig 3A, and 20°C GLMM, Estimate = 0.001, p = 0.035, Table 3SC Fig 3C*), this effect weakened at thermal extremes. This suggests that temperature modulates the relationship between host density and infection intensity (Fig 3). For example, at 23°C, as population size increased, burden decreased (*23°C GLMM, Estimate = −0.002, p = 0.024, Table 3SD Fig 3D*). Thus, the baseline temperature, timing in relation to the pulse event, and host population influence parasite proliferation. Yet, while population size influences parasite burden, it is also shaped by the combined effects of baseline temperature, timing relative to pulse events, treatment type, and parasite burden itself.

**Fig 3:**
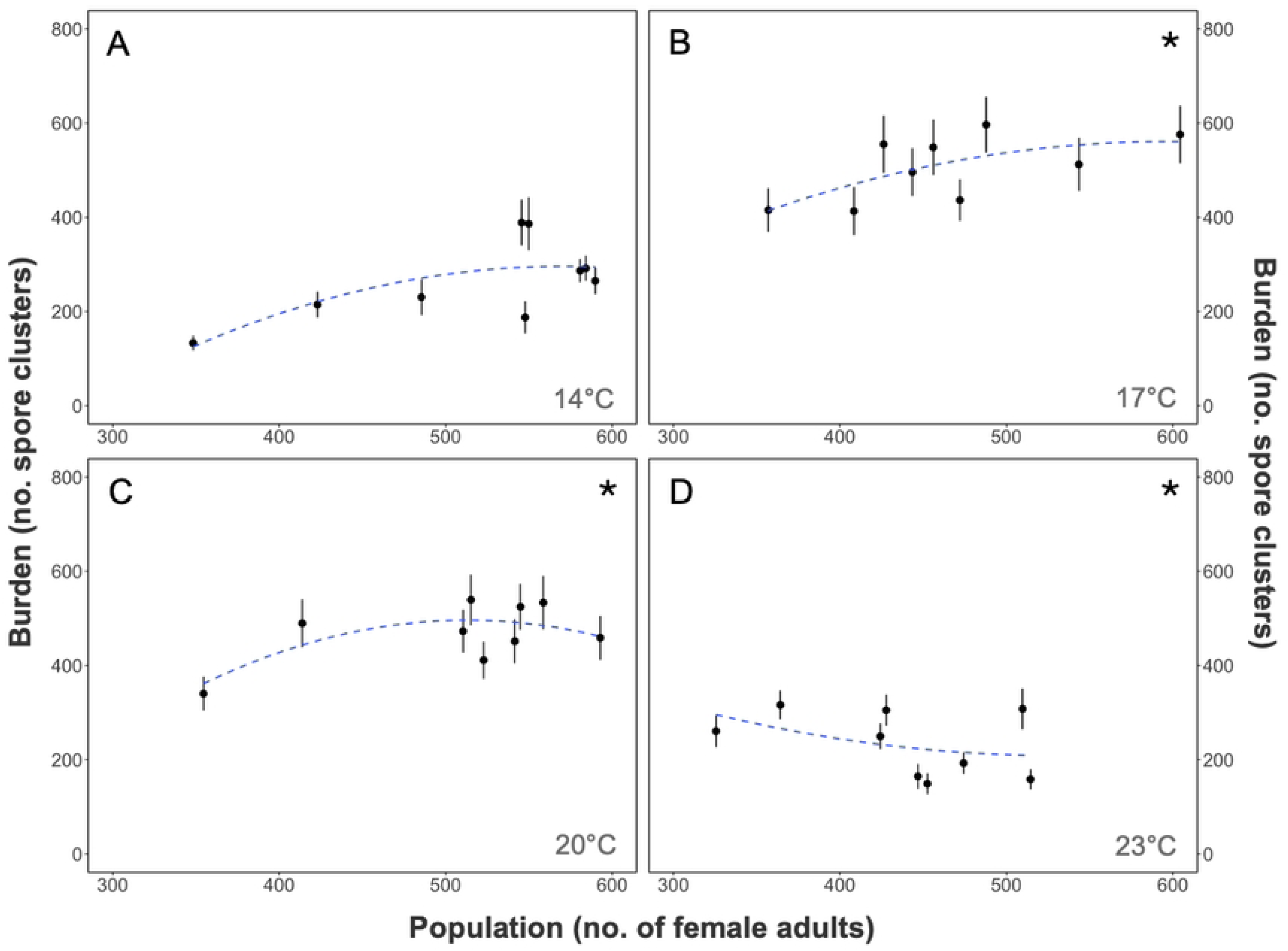
**The effect of population size on burden** in Daphnia magna infected with Ordospora colligata. Parasite burden (mean number of spore clusters, including zeros for uninfected but exposed hosts) and population size were averaged across nine weekly observations at four baseline temperatures: (A) 14°C, (B) 17°C, (C) 20°C, and (D) 23°C. Each spore cluster contains 32-62 individual Ordospora. Error bars represent the standard error, and each dashed line is the fit of the generalised linear mixed model (see Table 3S for statistical results). The presence of an asterisk (*) in a panel represents a significant relation between population size and burden.

### Population size

Like burden, host population size was influenced by baseline temperature, time in relation to the pulse event, treatment type (pulse or constant), and parasite burden. Population size, defined as the number of large reproducing adult females, was altered by up to 1.5-fold depending on the timing in relation to the pulse event and baseline temperature (Fig 3). For example, at 17°C the population size in the pulse treatment during the period ‘just after’ a heatwave occurred (week 6) was reduced by 182 *Daphnia* compared to constant temperature populations (Fig 3B) (*emmean, z(inf) = 4.829, p < 0.001, see Table 9S for p-values of all custom contrasts*). Moreover, at 17°C, all timings in relation to the pulse events resulted in differential outcomes in population size (*emmean, constant before vs pulse before, z(inf) = −4.548, p < 0.001, constant during vs pulse during, z(inf) = 4.179, p < 0.001, and constant after vs pulse after, z(inf) = 3.804, p < 0.001, see Table 9S*). While the burden increased with a heatwave at 14°C, the population size decreased in the periods ‘just after’ and ‘after’ the heatwave (*emmean, z(inf) = 2.763, p = 0.012, and z(inf) = 3.406, p = 0.002, respectively, Table 9S*). Although by the end of the experiment, the population size at 14°C had gradually increased to levels similar to those in the constant treatment (Fig 3A). Cold snaps had a lesser effect on population size, with a decrease in population size at 20°C but an increase in size at 23°C (Fig 3C, 3D) (*emmeans, constant during vs pulse during at 20*°*C, z(inf) = 2.379, p = 0.030, constant during vs pulse during at 23*°*C, z(inf) = −2.36, p = 0.030, Table 9S*). Overall, the timing in relation to the pulse event can influence population size mediated by baseline temperature (*GLMM, Type III Wald χ²_(12)_ = 88.41, p < 0.001, and Table 2*). Similarly, the effect of burden on population size varied with temperature, with marginal significance for an interaction (*GLMM, Type III Wald χ²_(3)_ = 7.84, p = 0.051; Table 2*). However, the negative association between population size and burden became stronger at higher temperatures (*GLMM, Estimate = –1.19, p = 0.023; Table 6S*)." Thus, all factors can influence the outcome of host population size.

**Table 2:** **Type III Wald chi-square test results for the generalised linear mixed model (GLMM) analysing population size** in Daphnia magna infected with Ordospora colligata. The Type III Wald χ² tests the significance of predictor variables, including temperature (modelled as a cubic polynomial), period, and population size. Significant values are bolded (See Table 7S for GLMM).

| Explanatory Variable | Chisq | Df | P value |
| --- | --- | --- | --- |
| (Intercept) | **117,340** | **1** | **<0.001** |
| Temperature | **59.747** | **3** | **<0.001** |
| Time Period | **27.219** | **4** | **<0.001** |
| Burden | 9.472 | 1 | 0.059 |
| Temperature: Time Period | **88.408** | **12** | **<0.001** |
| Temperature: Burden | **7.839** | **3** | **0.051** |

### Comparison of burden in populations and individuals

The population-level findings reflect similar patterns to those seen in individual-level experiments. The effect of a pulse treatment is dependent on baseline temperature at both population and individual level (Fig 4). For example, at 14°C, both population and individual burdens increase when a heatwave occurs (*emmean, z(inf) = 4.383, p < 0.001, and z(inf) = 2.527, p = 0.046, respectively, Table 3*). Trends were also similar at 17°C, whereby the burden increased in the heatwave treatment at both the population and individual levels (Fig 4). Similarly, a cold snap can reflect patterns seen at both the population and individual levels and is dependent on baseline temperature. For example, at 20°C, both the population and individual burden decreased when a cold snap occurred compared (Fig 4). Therefore, individual-level exposure patterns are reflected at the population level, with treatment effects varying depending on baseline temperature.

**Fig 4:**
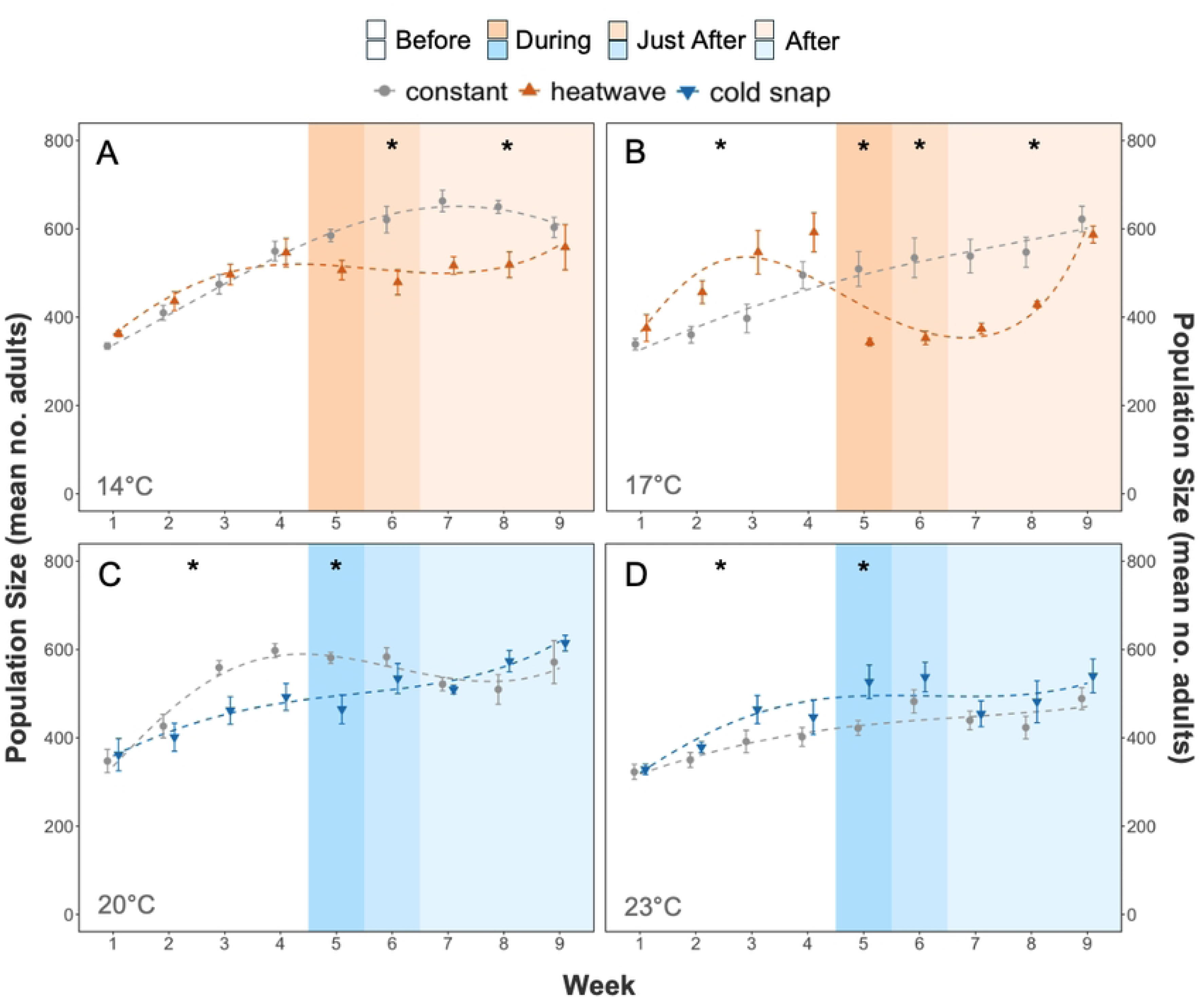
**The temporal effects of pulse temperature treatments on population size** in Daphnia magna infected with Ordospora colligata. Population size (mean number of adults) is shown over the nine recorded weeks at four baseline temperatures: (A) 14°C, (B) 17°C, (C) 20°C, and (D) 23°C. Each spore cluster contains 32-62 individual Ordospora. Error bars represent the standard error, and each dashed line is the cubic fit of the generalised linear mixed model (see Table 6S for statistical results). Grey; constant treatment, orange; heatwave pulse treatment, blue; cold snap pulse treatment. Shaded regions mark experimental periods: Before (white), During (orange/blue), Just After (lighter orange/blue), and After (lightest orange/blue). Asterisks (*) indicate significant differences in burden between pulse and constant treatments in specific time periods (emmeans contrasts, Table 9S).

**Fig 5:**
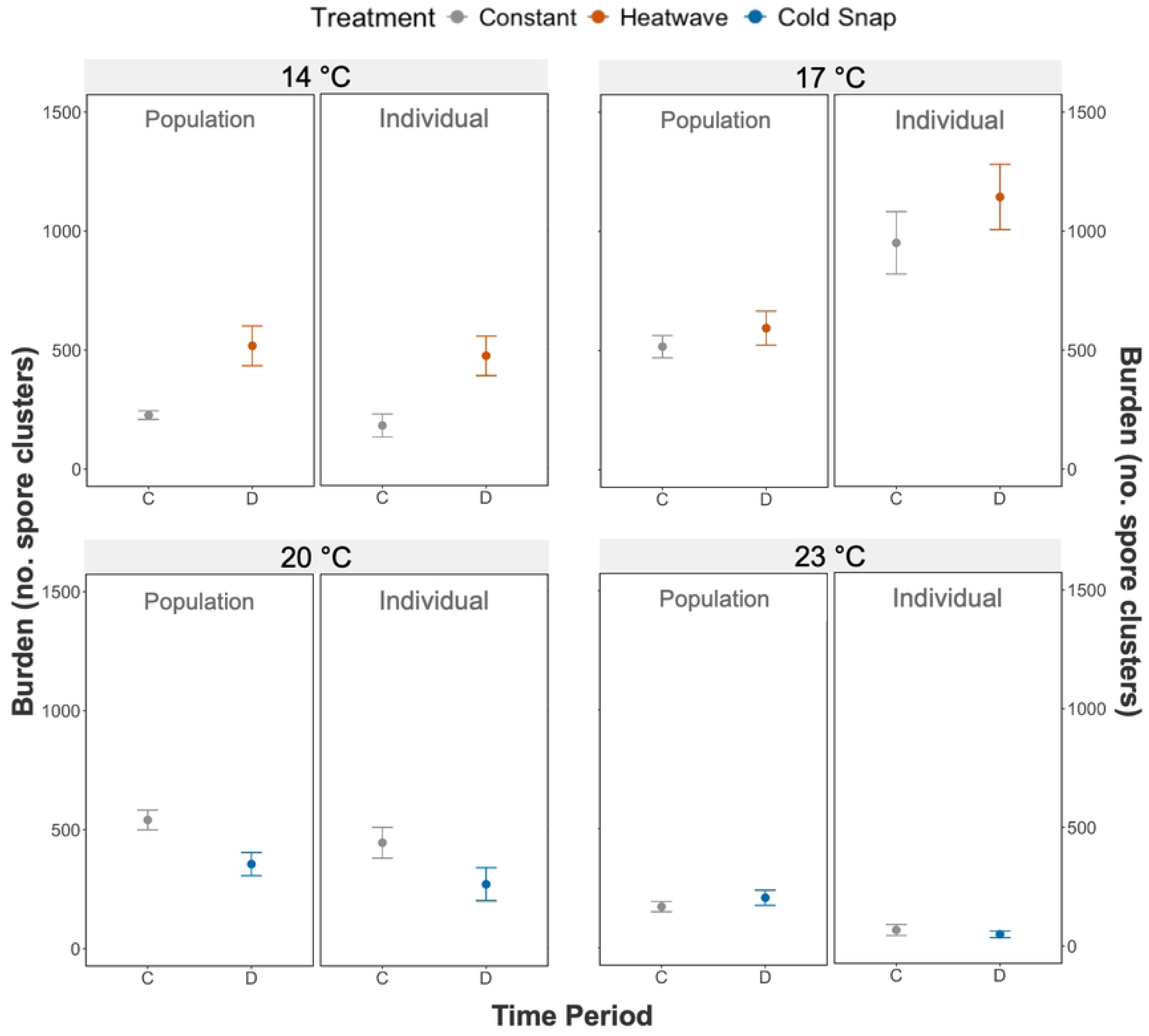
**Comparison between the effects of pulse and constant treatments on burden at both individual and population scales** in Daphnia magna infected with Ordospora colligata. Burden (number of spore clusters, including zeros for uninfected but exposed hosts) is shown over the nine recorded weeks at four baseline temperatures: 14°C, 17°C, 20°C, and 23°C. Each spore cluster contains 32-62 individual Ordospora. Error bars represent the standard error. Grey; constant treatment, orange; heatwave pulse treatment, blue; cold snap pulse treatment. C; constant (no pulse treatment), D; during pulse period.

**Table 3:** **Custom contrasts for burden comparison between periods at both population and individual levels** in Daphnia magna infected with Ordospora colligata. Each pulse treatment period was compared to its constant equivalent at the same temperature or to the ‘before’ pulse period. The response variable was burden, with uninfected but exposed hosts coded as zero. The ‘Inf’ degrees of freedom reflect the model’s complexity and the estimation process. Exp; experiment, CS; cold snap individual experiment, HW; heatwave individual experiment, POP; population experiment. Significant contrasts are bolded

| Exp | Temp | Contrast | Estimate | SE | Df | z ratio | p value |
| --- | --- | --- | --- | --- | --- | --- | --- |
| **POP** | **14** | **during HW vs constant** | **0.827** | **0.189** | Inf | **4.383** | **0.0001** |
| **HW** | **14** | **during HW vs constant** | **0.875** | **0.346** | Inf | **2.527** | **0.046** |
| POP | 17 | during HW vs constant | 0.117 | 0.189 | Inf | 0.621 | 0.5544 |
| HW | 17 | during HW vs constant | 0.185 | 0.312 | Inf | 0.591 | 0.5544 |
| POP | 20 | during CS vs constant | -0.431 | 0.189 | Inf | -2.282 | 0.0599 |
| CS | 20 | during CS vs constant | -0.431 | 0.321 | Inf | -1.345 | 0.3571 |
| POP | 23 | during CS vs constant | 0.174 | 0.189 | Inf | 0.919 | 0.4773 |
| CS | 23 | during CS vs constant | -0.337 | 0.357 | Inf | -0.944 | 0.4773 |

## Discussion

The effect of heatwaves and cold snaps on disease dynamics and parasite fitness is complex and context-dependent. Temperature fluctuations influence both host population size and parasite burden, often in nonlinear and unpredictable ways. Understanding these dynamics is crucial, as temperature-induced changes in infection levels and host population size can have cascading effects on ecological processes and disease transmission [35].

### Burden and population size

Parasite burden and population size were not independent; rather, their relationship varied with baseline temperature, consistent with interactive effects of ecological context on disease dynamics. Larger host populations generally supported higher numbers of pathogens, consistent with density-dependent transmission dynamics [36], where increased host density facilitates more frequent contact between infected and susceptible individuals. This pattern has been observed across many systems, for example, amphibian-chytrid fungus [37], guppy-platyhelminth [38], and moth-fungus [39] interactions. However, at higher temperatures (23 °C), this positive relationship reversed, with larger populations linked to lower burdens. This decoupling suggests that high temperatures may disrupt transmission dynamics, resulting in decreased burdens at higher densities. Such reversals are consistent with negative density dependence, where increased host density can suppress, rather than amplify, infection under certain ecological conditions [40]. Several mechanisms could contribute to this reversal, for example, reduced parasite performance [13] or decreased parasite viability [41]. Moreover, crowding at high temperatures could intensify competition for food or space, leading to behavioural or physiological changes [42, 43], and enhanced host immunity [44, 45], resulting in reduced transmission efficiency. Additionally, high-density conditions may promote avoidance behaviours, like reduced aggregation or increased movement, further limiting spore encounter rates [46]. Similar temperature-driven interference with transmission has been observed in *Daphnia dentifera* infected with *Metschnikowia bicuspidata,* where foraging interference and high host density at high temperatures decreased parasite invasion and prevalence [47]. Moreover, the negative relation between density and burden at 23°C may reflect that animals experiencing both thermal and crowding stress are unable to withstand higher parasite burdens, resulting in higher host mortality rates [25]. For example, in three-spined sticklebacks (*Gasterosteus aculeatus*), fish with higher parasite loads experienced increased mortality rates under elevated temperatures [48]. Together, these findings suggest that the relationship between host population size and burden is modulated by ecological context and can shift nonlinearly, with extreme temperatures potentially disrupting density-dependent transmission and amplifying disease-induced population decline. This complexity underscores the need to consider feedback between host demography and parasitism in predicting responses to climate variability.

### Effect of cold snaps

Cold snaps can increase the burden depending on the baseline temperature. For example, at 23°C, a cold snap was followed by a delayed increase in burden, without a corresponding change in population size. Indeed, a cold snap in warmer climates can sharply increase fungal disease risk in wildlife populations [49]. When the thermal optima of the host and parasite differ, fluctuations in temperature can shift the balance in their interaction, resulting in either host or parasite having an advantage over the other, as stated by the Thermal Mismatch Hypothesis [32]. In other systems, for example, the Red-backed Salamander (*Plethodon cinereus*) and the chytrid fungus (*Batrachochytrium dendrobatidis*), warm-adapted hosts had a higher probability of infection at cool temperatures compared to cold-adapted hosts [50]. Furthermore, temperature changes may compromise host physiology by suppressing immune responses or slowing metabolic activity, temporarily reducing the host’s ability to resist infection [8]. For example, cold stress compromised the immune function of the Madagascar hissing cockroach (*Gromphadorhina coquereliana*) [51]. Here, the suppression of *Daphnia* immunity during the cold snap and subsequent rewarming could have contributed to the delayed increase in burden at higher temperatures [52]. An increase in burden was also previously noted at the individual level by McCartan et al. (2024), where burden was increased after a cold snap occurred at 23°C in the same host-parasite system. Thus, cold snaps can shift the outcome of the *Daphnia–Ordospora* interaction in a temperature-dependent manner, highlighting the close relationship between infection intensity and host population dynamics.

### Effect of heatwaves

Heatwaves, like cold snaps, can alter disease in the *Daphnia-Ordospora* system in a temperature-dependent way. Indeed, an increase in burden following a heatwave was observed at 14°C. Here, the rapid temperature rise may have overwhelmed the capacity of the *Daphnia* to maintain homeostasis, leading to a decrease in population size. In contrast, *Ordospora* may have acclimatised faster to the heatwave, increasing replication and spore shedding, thereby raising exposure risk and leading to a higher burden during and after the heatwave occurred. Similar increases in burden near the lower end of the thermal extreme are also present in other systems [53, 54]. For example, daily temperature fluctuations are expected to increase malaria transmission at lower baseline temperatures while decreasing transmission at higher temperatures [55]. Increases in burden following thermal fluctuations can be explained by the Thermal Variability Hypothesis [31]. This theory posits that, as parasites are generally smaller than their hosts and have higher metabolic rates (according to the Metabolic Theory of Ecology [56]), they can physiologically adjust more quickly to temperature changes, giving them a temporary advantage. This is supported by other studies using the same host-parasite system, which found that short-term fluctuations led to an increase in burden [24, 57]. Our results furthermore show that such increases can persist up to four weeks after a heatwave (at 14°C), indicating that parasites may derive long-term fitness benefits from short-term extreme weather events.

Long-term increases in disease burden following extreme weather conditions may arise from sustained changes in host physiology [58], immune function [59], or shifts in the host-parasite dynamics that are not quickly reversed once temperatures stabilise [60]. For example, increased air temperatures following heatwaves elevated *Perkinsus marinus* (a protozoan parasite) infection intensity in oyster populations and suppressed immune defences, leading to prolonged infection levels even after temperatures normalised [61]. These findings raise important questions about the broader consequences of more frequent extreme weather events. For instance, if parasites continue to benefit disproportionately from transient temperature spikes, this could extend the duration of outbreaks, shift their seasonal timing, or amplify their severity [62, 63]. Thus, brief thermal events can trigger prolonged shifts in disease dynamics, potentially altering the trajectory of epidemics and influencing long-term host population health. As climate change increases the frequency and intensity of heatwaves, understanding these lasting effects is essential for predicting parasite persistence and outbreak potential under future environmental scenarios, especially as heatwaves at 17°C did not cause the same effect.

There was no observed increase in spore burden at 17°C, despite reductions in population size both during and after the event. While we observed no response, potential opposing effects of heatwaves on both host and parasite life history traits may have masked any changes in burden. Given the narrower thermal tolerance for *Ordospora* (11.8-29.7°C) compared to its *Daphnia* host (6-33.3°C) (41), the heatwave could have imposed physiological stress on the parasite (Thermal Stress Hypothesis [31]). However, this may be offset by increased contact rates with higher temperatures due to increased filtration rates of the host [64]. Other systems have shown that different effects on host and parasite traits can cancel each other out [65]. A similar lack of response to temperature variation has been observed in other systems, such as the monarch butterfly and its protozoan parasite *Ophryocystis elektroscirrha*, where both host and parasite show reduced performance under heat stress [25]. Thus, while long-term effects of a heatwave on parasite burden were observed at 14°C, no effects were present at 17°C, highlighting that the effect of a heatwave, similar to cold snaps, can depend on the mean temperature at which the extreme weather event occurs.

### Population versus individual level

When pulse treatment effects were compared across levels, specifically, individual-level experiments where parasites were exposed to long, strong pulses during infection, and population-level experiments where burden was measured during the pulse itself, there was strong concordance in responses. For example, the observed increase in burden during a heatwave at 14°C and a decrease in burden during a cold snap at 20°C were consistent across both scales. This suggests that temperature-driven effects on parasite fitness may scale predictably from individuals to populations. Thus, individual-level physiological responses, such as temperature-induced changes in host immunity [8] or parasite replication [66], may translate into population-level infection dynamics. For instance, temperature dependence around a thermal optimum can be scaled from the individual to the population level, although not always proportionally [67]. That is, while rising mean temperatures may consistently affect burden at the individual level, the population-level outcomes (such as prevalence or outbreak size) may also be shaped by ecological complexity [68]. For example, in a snail-schistosome system, individual-level traits like body size influenced host exposure and susceptibility, leading to predictable patterns of infection prevalence at the population level [69]. Host density [70], resource competition [71], and environmental carrying capacity [72] may modulate the strength and direction of temperature effects at different biological levels. Thus, integrating both physiological and ecological perspectives is essential to fully understand how temperature variability shapes disease dynamics across biological scales.

### Conclusion

In conclusion, the effect of extreme temperature variation on parasite fitness is complex. It depends on the baseline temperature at which it occurs and the type of variation, with a heatwave having long-lasting effects (>4 weeks) on burden. Host population size also exhibited temperature-dependent patterns, highlighting the broader effects of environmental variation on demographic dynamics. Importantly, parasite burden and host density were interdependent; each influenced the other in ways that shaped infection dynamics. At lower temperatures, burden and population size had a positive relationship; however, at higher temperatures, higher host densities resulted in lower burdens. Trends in burden were broadly consistent across both population- and individual-level, highlighting the potential for individual physiological responses to scale up and influence population-level disease outcomes. These findings underscore the importance of considering both environmental context and temporal dynamics when assessing host-parasite interactions under climate change. As climate change is resulting in increased frequency and intensity of anomalous weather events [73], understanding the influence of these on host-parasite dynamics will become all the more important. Given the key role of *Daphnia* in their ecosystem, the effect can have far-reaching impacts [74]. For example, a reduction in *Daphnia* density due to infection by another parasite led to alterations in the food web, causing a trophic cascade [75]. Moreover, accounting for temperature variation in predictive models will be essential for more robust predictions [76], as parasites can alter co-evolutionary dynamics [77], reshape ecological networks [78], and drive population-level changes in host species [79]. Failing to consider temperature fluctuations may lead to overestimates in parasite fitness traits at high temperatures [80]. Thus, identifying underlying mechanisms will be essential for the accurate assessment of natural populations in the face of climate change.

## Materials and Methods

### Study system

*Daphnia magna* is a freshwater crustacean found across the northern hemisphere and is a key species in many aquatic ecosystems [74]. This is a popular model system for environmentally transmitted diseases (characterised by horizontal transmission mechanisms and infection dynamics governed by mass action principles) [81]; for example, due to their fast generation times, well-known ecology [82], and parthenogenic life cycle [83]. *Ordospora colligata* is a microsporidian microparasite infecting many genotypes of *D. magna* and is horizontally transmitted to the host via filter feeding [84]. The parasite invades the upper gut epithelium, forming intracellular clusters of 32-64 spores, which burst and shed into the water via faeces [83]. Moreover, *Daphnia* have a wider thermal range of 6-33.3°C compared to *Ordospora’s* narrower range of 11.8-29.7°C [85], which may cause temperature shifts that disproportionally affect one of the antagonists. Here we used *Daphnia magna* genotype Fi-OER-3-3 and *Ordospora colligata* isolate 3, originally sampled from Tvärminne, Finland, and both have been maintained in the laboratory cultures for at least 10 years.

### Experiment design

To determine whether extreme weather can alter disease dynamics in the *Daphnia-Ordospora* system, we used a factorial experimental design, testing two temperature regimes (constant and pulse) at four baseline temperatures (14, 17, 20, and 23°C). All pulse treatments occurred eight weeks after the experiment began (that is, four weeks after measurements started) (Fig 1). Temperature pulses either consisted of a heatwave or a cold snap, depending on the baseline temperature, with lower temperatures experiencing a heatwave (14 and 17°C) and higher temperatures a cold snap treatment (20 and 23°C). A heatwave treatment increased the temperature by +6°C for 10 days, while a cold snap decreased by − 6°C for 10 days relative to the baseline temperature. Once the pulse concluded, the treatment was returned to its original baseline temperature. In total, there were eight treatments (two temperature regimes • four baseline temperatures), each with five replicates, for a total of 40 populations (Fig 1). Every replicate started with a population of 300 *Daphnia,* kept in a 5 L mesocosm (filled with 2 L of Artificial *Daphnia* Medium (ADaM [86]), in which infection was well established. Temperature was manipulated by placing each of the mesocosms into temperature-controlled water baths. There were three baths per baseline temperature, and treatments were spread between baths. Of these, two baths per baseline temperature contained two constant and two pulse treatments, and the remaining bath per baseline temperature contained one constant and one pulse treatment.

### Temperature-controlled water baths

In total, there were twelve temperature-controlled water baths (three baths per baseline temperature). Each bath was equipped with an aquarium chiller (Hailea HC-150A, DC300, or DC750), a heater (EHEIM JÄGER 300W), pumps (Micro-Jet Oxy and Oase Optimax 500) and a programmable controller (Inkbird ITC-308 or ITC-310T) which regulated the temperature. Temperatures in each bath were continuously monitored in an extra mesocosm filled with only ADaM using a HOBO logger (HOBO UA-001-08) and checked daily. While pumps ensure a uniform temperature distribution, mesocosms were also rotated clockwise daily to negate any temperature variation within the baths. To simulate temperature pulses (i.e., heatwaves or cold snaps), population mesocosms were moved between designated baths, while the baths were maintained at a constant baseline temperature for the duration of the experiment. After moving mesocosms, they reached the target temperatures within an hour. After ten days, the pulse period ended, and the mesocosms were returned to their original positions.

### Daphnia preparation

To initiate the experiment, twelve large populations of *Daphnia* (∼12,000 animals) exposed to *Ordospora* were set up twelve weeks prior to the beginning of the experiment using the following method. These populations were generated from existing stock populations that were kept in the laboratory for over five years. The existing stock populations were filtered using a coarse (1 mm) mesh filter, and animals were added to a fresh 5 L mesocosm with ∼2 L of fresh Artificial *Daphnia* Medium (ADaM) [86] (Fig 1). To set up twelve mesocosms, small amounts from each stock population were added to the empty mesocosms until all housed equal volumes of ADaM and animals. Each of the mesocosms was fed *ad libitum* with batch culture algae (*Scenedesmus* sp.) [87]. These populations were maintained by separating accumulated debris from the adult animals every four weeks using a coarse (1 mm) mesh filter. Animals caught by the filter were transferred to a clean mesocosm with 500 mL of the original ADaM (to retain juveniles and *Ordospora* spores), and 1500 mL of fresh ADaM was added to replenish the medium.

After the final maintenance, right before the beginning of the experiment, five animals in each mesocosm were checked for infection, and the presence of the pathogen was verified for all twelve mesocosms. Then, all animals from the twelve mesocosms were pooled and mixed to ensure uniform exposure across populations before redistribution. To set up each of the 40 experimental populations, 300 *Daphnia* of different developmental stages (e.g. juveniles and adults) were added to a 5 L mesocosm with 2 L of ADaM and fed 20 mL of 18 million algae/mL. The replicates were staggered, so all replicate-one populations began on day 1 (i.e., eight populations), then all replicate-two populations began on day 2, and so on. Populations were maintained for four weeks after initialisation to stabilise before measurements of burden and population size were taken.

### Population Maintenance

Populations were maintained every seven days. Before filtering a population, any large debris was removed from the mesocosm manually by using a pipette. The population was sifted twice through a 1320 µm mesh to catch all adults and larger debris; the remaining medium was then sifted through a fine mesh (200 µm) to catch the juveniles. Fifteen per cent of the original medium was replaced with fresh ADaM (i.e., 300 mL of fresh ADaM, 1700 mL of original ADaM) (Fig 1). Around half of this medium was then used to gently wash the adult *Daphnia* away from the mesh into a temporary mesocosm while keeping the remaining debris caught in the mesh. The other half of the medium was used to wash the fine mesh filter into a separate mesocosm. This enabled independent recording of the adult population size. When measurements were completed, the adult *Daphnia* were added back in with the remaining population. Feeding occurred three times a week when mesocosms received 20 mL of 18 million algae/mL.

### Measurement of population size and parasite fitness

Measurements of population size and parasite fitness were conducted weekly after populations were stabilised. Immediately after filtration over the 1320 µm mesh during maintenance, the population sizes of the adults were recorded to assess the number of large females in a population. A Canon 2000D camera was attached to a tripod and was hung with the lens 15 cm above a 2 L semi-transparent container. A flat light box was placed below the semi-transparent container to illuminate the individuals from below to increase contrast. The adult *Daphnia* in the temporary mesocosm were added to this semi-transparent container, and the camera zoom was altered so it filled the frame of the camera (Fig 1). Two videos were recorded for each set of adult *Daphnia*, all lasting four seconds. Once the videos were taken, both the adults and the remaining *Daphnia* were added to a single mesocosm and placed back into the water bath. All videos were analysed for population counts using the R package ‘trackdem’ [88]. This package works by subtracting moving particles from the background, which can then be counted when appropriate thresholds are set. The package ‘trackdem’ uses images captured from a video, which were extracted using the application ‘ffmpeg’ in the computer terminal. Two frames per second were captured from the four-second videos. The channel used was blue, and the average threshold used was −0.12. Any remaining *Daphnia* that the software did not detect were manually counted to ensure the most accurate population count was recorded.

On the same day population size was quantified, six adult *Daphnia* of varying sizes per mesocosm were sampled to measure the infection prevalence (i.e., presence or absence of spores) and burden (i.e., the number of spore clusters in the host) of *O. colligata*. If infected, spore clusters (each cluster holding up to 64 individual spores) were counted using bright field or phase-contrast microscopy (with 400x magnification). Any males (n = 1) or inconclusive infections (n = 6) were omitted from the analysis. After nine timepoints (weeks) of data collection, the experiment concluded.

### Data analysis

Analysis was performed using R version 2024.04.2+764 [89]. The response variables were burden (i.e., the number of *Ordospora* spore clusters, including zeros for uninfected exposed individuals) and population size (i.e., the mean count of adults from population videos). For both models, ‘period’ was a factor which included the different time points in the experiment: constant treatment, before pulse (weeks 1-4), during pulse (week 5), just after pulse (week 6), and after pulse (weeks 7-9). This structure allowed for an extended Before-After-Control-Impact (BACI) design to compare temporal responses between pulse and constant treatments [90]. ‘Mesocosm’ was included as a mixed effect variable to account for batch-specific variability; the remaining variables were fixed effects. Therefore, when analysing burden, the predictor variables were ‘baseline temperature’, ‘period’, ‘population size’, and ‘mesocosm’. When analysing population size, the predictor variables were ‘baseline temperature’, ‘period’, ‘burden’, and ‘mesocosm’. For the burden model, ‘baseline temperature’ had a quadratic polynomial to accommodate the non-linear temperature response, while in the population size model, it had a cubic relationship. To avoid issues with convergence and collinearity in both models, ‘baseline temperature’ was centred. Moreover, in the burden model, ‘population size’ was centred, and in the population size model, ‘burden’ was scaled. A Generalised Linear Mixed Model (GLMM) with a negative binomial link function was fitted using the ‘glmmTMB’ package [91] to account for overdispersion in count data and batch-specific effects. An analysis of deviance (type III Wald chi-square test) was then used to investigate the relationship between variables and assess the overall significance of the predictor variables. Custom contrasts were created with the “emmeans” package [92] to compare treatments in both burden and population size. For this, ‘baseline temperature’ was treated as a factor and modelled with ‘period’ as an interaction. The ‘constant’ period was split up according to the pulse treatments for discrete comparisons, (i.e., before constant (weeks 1-4), during constant (week 5), just after constant (week 6), and after constant (week 7-9)), and treatments were compared accordingly. The contrast p-values were adjusted using the ‘Benjamini-Hochberg’ method for multiple comparisons [93].

The population-level results were then compared to individual-level findings from previous papers by McCartan et al. (2024) and McCartan et al. (2025). Data which was comparable to the population-level results was extrapolated from both data sets. Specifically, for heatwave data, the data taken was from the baseline temperatures 14 and 17°C, where constant temperature data and heatwave treatments occurred on day 0 (day of infection), and lasted for 6 days with an amplitude of +6°C. For cold snaps, the data taken was from the baseline temperatures of 20 and 23°C. Here, constant temperature data was also used, and the cold snaps lasted for 6 days with an amplitude of −6°C. A custom contrast was then created, where pulse and constant treatments were compared to each other within the same experiment. The contrast p-values were adjusted using the ‘Benjamini-Hochberg’ method for multiple comparisons.

## Acknowledgements

The authors thank Dieter Ebert and Jürgen Hottinger for the provision of the biological materials, Alison Boyce and Sinead Kelly for technical assistance in creating the water baths and maintenance of electrical equipment, and Whitney Parker for helping with empirical work.

## Supporting Information Captions

***Table 1S: Generalised Linear Mixed Model (GLMM) output for burden** in Daphnia magna infected with Ordospora colligata. Temperature was modelled as a quadratic polynomial to account for non-linearity and is presented separately by degree (linear [L] and quadratic [Q] The GLMM included fixed effects of temperature (modelled as a quadratic polynomial), time period (relative to a heatwave or cold snap), and population size, with a random effect of bucket. The response variable was spore burden, with uninfected but exposed hosts coded as zero. Significant values are bolded*.

***Table 2S: Type III Wald chi-square test results for the generalised linear mixed model (GLMM) analysing burden** in Daphnia magna infected with Ordospora colligata. The analysis of deviance tests the significance of predictor variables, including temperature (modelled as a quadratic polynomial), period, and population size. The response variable was spore burden, with uninfected but exposed hosts coded as zero. Significant values are bolded*.

***Table 3S: Temperature-specific Generalised linear mixed model (GLMM) outputs for burden** in Daphnia magna infected with Ordospora colligata. A; For 14°C, B; for 17°C, C; for 20°C, D; for 23°C. The GLMM included fixed effects of time period (relative to a heatwave or cold snap), and population size, with a random effect of bucket. The response variable was spore burden, with uninfected but exposed hosts coded as zero. Significant values are bolded*.

***Table 4S: Estimated marginal means (emmeans) for burden** in Daphnia magna infected with Ordospora colligata. The estimates were derived using a generalised linear model (GLM) with baseline temperature and treatment period as factors, using a negative binomial family. The response variable was spore burden, with uninfected but exposed hosts coded as zero. Values are presented on the log scale for appropriate interpretation of the data. The ‘Inf’ degrees of freedom reflect the large sample size and the flexibility of the model*.

***Table 5S: Custom contrasts for burden ‘emmeans’ comparing treatments across baseline temperatures and periods** in Daphnia magna infected with Ordospora colligata. Each pulse treatment period was compared to its constant equivalent at the same temperature or to the ‘before’ pulse period. The response variable was spore burden, with uninfected but exposed hosts coded as zero. The ‘Inf’ degrees of freedom reflect the model’s complexity and the estimation process. C; constant treatment, P; pulse treatment. Significant contrasts are bolded*.

***Table 6S: Generalised Linear Mixed Model (GLMM) output for population size** in Daphnia magna infected with Ordospora colligata. Temperature was modelled as a cubic polynomial to account for non-linearity and is presented separately by degree (linear [L] and quadratic [Q], and cubic [C]). A random intercept for bucket was included to account for repeated measures. Significant values are bolded*.

***Table 7S: Type III Wald chi-square test results for the generalised linear mixed model (GLMM) analysing population size** in Daphnia magna infected with Ordospora colligata. The analysis of deviance tests the significance of predictor variables, including temperature (modelled as a cubic polynomial), period, and population size. Significant values are bolded*.

***Table 8S: Estimated marginal means (emmeans) for population size** in Daphnia magna infected with Ordospora colligata. The estimates were derived using a generalised linear model (GLM) with baseline temperature and treatment period as factors, using a negative binomial family. Values are presented on the log scale for appropriate interpretation of the data. The ‘Inf’ degrees of freedom reflect the large sample size and the flexibility of the model*.

***Table 9S: Custom contrasts for population size ‘emmeans’ comparing treatments across baseline temperatures and periods** in Daphnia magna infected with Ordospora colligata. Each pulse treatment period was compared to its constant equivalent at the same temperature or to the ‘before’ pulse period. The response variable was spore burden, with uninfected but exposed hosts coded as zero. The ‘Inf’ degrees of freedom reflect the model’s complexity and the estimation process. C; constant treatment, P; pulse treatment. Significant contrasts are bolded*.

***Table 10S: Estimated marginal means (emmeans) for burden comparison between periods in both population and individual experiments** in Daphnia magna infected with Ordospora colligata. The estimates were derived using a Generalised Linear Model (GLM) with treatment period as a factor, using a negative binomial family. Treatment combined the factors: experiment, temperature, treatment, and period. The response variable was spore burden, with uninfected but exposed hosts coded as zero. Exp; experiment, CS; cold snap individual experiment, HW; heatwave individual experiment, POP; population experiment. Period; constant; continuous baseline with no pulse, during; the week when the pulse occurred. Values are presented on the log scale for appropriate interpretation of the data. The ‘Inf’ degrees of freedom reflect the large sample size and the flexibility of the model*.

***Table 11S: Custom contrasts for burden comparison between periods in both population and individual experiments** in Daphnia magna infected with Ordospora colligata. Each pulse treatment period was compared to its constant equivalent at the same temperature or to the ‘before’ pulse period. The response variable was spore burden, with uninfected but exposed hosts coded as zero. The ‘Inf’ degrees of freedom reflect the model’s complexity and the estimation process. Exp; experiment, CS; cold snap individual experiment, HW; heatwave individual experiment, POP; population experiment*.

## Notes

### Competing Interest Statement

The authors have declared no competing interest.

